# Syndecan 2 regulates hematopoietic lineages and infection resolution in zebrafish

**DOI:** 10.1101/2020.05.04.076786

**Authors:** Bhawika Sharma Lamichhane, Brent W. Bisgrove, Yi-Chu Su, Bradley L. Demarest, H. Joseph Yost

**Author notes:** Corresponding author: H. Joseph Yost, University of Utah Molecular Medicine Program, Eccles Institute of Human Genetics, Bldg. 533, Room 3160, 15 North 2030 East, Salt Lake City, UT 84112-5330.

## Abstract

Syndecan 2 (*Sdc2*) is a transmembrane cell-surface heparan sulfate proteoglycan (HSPG) that has been implicated in the regulation of cell-cell signaling pathways and cell-matrix interactions. Surprisingly, homozygous recessive maternal zygotic (MZ) *sdc2* null mutants in zebrafish appear to have normal development, normal morphology and are viable and fertile in adulthood. Whole transcriptome RNA sequencing, FACS analyses, and imaging of transgenic reporter lines that distinguish specific hematopoietic lineages revealed that *sdc2* mutants have defects in the specification and proportions of red blood cells and neutrophils that initiate during embryonic hematopoiesis and likely persist through adulthood. During bacterial infections, MZ*sdc2* mutants have markedly reduced neutrophil recruitment and significantly higher death rates. Hematopoietic stem/progenitor cell (HSPC) numbers are also significantly reduced in MZ*sdc2* mutants. In zebrafish, cells that bud off of the ventral region of somites are thought to give rise to the reticular stromal cells of the caudal hematopoietic tissue (CHT) stem cell niche. In MZ*sdc2* mutants, these budding cells have abberant blebbing morphology associated with widespread apoptosis during induction of HSPCs and with changes in the vascularization and stromal cell structure of the CHT stem cell niche. This suggests that loss of *sdc2* disrupts the earliest events of definitive hematopoiesis. Our findings of hematopoietic defects, nascent immune system alterations and inability to resolve infection in *sdc2* mutants sets the stage for examining the roles of HSPG genes in a wide range of hematopoietic and immune defects in humans.

**Key Points:**

1. Syndecan 2 regulates the formation of Hematopoietic Stem/Progenitor Cells, differentiation into hematopoietic populations and the CHT architecture
2. Syndecan 2 mutants are significantly more susceptible to bacterial infection

## Introduction

Syndecans are transmembrane proteoglycans that have been proposed to modulate ligand-dependent activation of major signaling pathways to facilitate cellular processes and regulate extracellular matrix (ECM) architecture during development and disease. The glycosaminoglycan chains on syndecans facilitate direct binding with growth factors, cytokines, chemokines, and ECM proteins to allow for receptor docking, concentration and protection of molecules at cell surfaces, thereby modulating intracellular signaling^1^. The cytoplasmic domain of syndecans, with binding sites for cytoskeletal proteins, promotes focal adhesion formation, matrix assembly and remodeling^2–4^. Syndecans, along with integrins, serve as a scaffold that connects the actin cytoskeleton to ECM proteins like fibronectin. This connection is essential for cell viability, and the same protein machinery is manipulated during cellular processes such as migration, adhesion, differentiation and anoikis.

There are three syndecan genes in zebrafish, *syndecan 2 (sdc2), syndecan 3 (sdc3)* and *syndecan 4* (*sdc4*)^5^. Syntenic analysis between zebrafish, mouse and humans has shown that zebrafish *syndecan 2* is the ortholog of human and mouse syndecan 2 and clusters to the vertebrate *sdc2* family^6^. Many interesting developmental phenotypes for *sdc2* came out of morpholino and dominant-negative overexpression studies in vertebrate models, but functional studies using knockout approaches have been sparse with only one study in mouse showing angeiogenic and arteriogenic defects in global and endothelial specific *Sdc2* knockouts.^7–11^

To further understand and explore the function of *sdc2*, and given concerns about non-genetic morpholino and dominant negative approaches^12,13^, we created *sdc2* mutants in zebrafish. Both the zygotic and maternal zygotic (MZ) mutants had no apparent phenotypes. Given that model organisms housed in artificial environment might not reveal gene expression changes as apparent phenotypes, we used whole-genome paired end RNA sequencing (RNA-seq) on MZ *sdc2* mutants and WT embryos at several stages of early development to observe if any genes regulating key biological processes were altered.

Differential gene expression analysis suggested that MZ*sdc2* mutants had alterations in hematological pathways. We found that *MZsdc2* mutants have defects in some of the earliest steps of definitive hematopoiesis, during the specification of HSPCs and stromal cells in the CHT niche.^14^ We uncover, for the first time, the role of *Sdc2* in the specification of hematopoietic cell populations which leads to aberrant immune response and failure to survive bacterial infection.

## Methods

Extended Materials and Methods are in supplemental material.

Creation of *sdc2* mutants: TALENs were designed against *sdc2* exon 2 using TALE-NT (http://boglabx.plp.iastate.edu/TALENT/TALENT/#). Site targeted in exon 2 was: <u>CCACGACAGAtGACCTGT</u>ACCTGGAGGAGGCTGG<u>ATCTGGAGGATACCCTGAAG</u>

DNA sequences corresponding to left and right TALENs were cloned into pCS2 vector (pCS2TAL3-sdc2TALEN-DDD, pCS2TAL3-sdc2TALEN-RRR) at University of Utah Mutation Generation and Detection Core. RNA encoding left and right TALENs were transcribed from Not1 digested vector using SP6 mMessage mMachine kit (Thermofisher). 50 pg RNA was injected into AB zebrafish embryos at 1-2 cell stage. Mutation efficiencies in injected G0s were determined by High-Resolution Melt Analysis (HRMA) and sibling embryos raised to adulthood. Adult G0s were outcrossed to wild-type AB line and embryos scored for the mutation by HRMA. Sibling embryos from clutches containing *sdc2* mutants were raised to adulthood and propagated. Present studies were carried out using animals homozygous for the *sdc2^zy^*^37^ mutation, a 7 bp deletion.

RNA-seq analysis: Individual RNA-seq samples were prepared from 30 wild-type and MZ*sdc2* mutants embryos at 5 stages: 1 somite, 16 somites, 30 hours post-fertilization (hpf), 48 hpf, and 72 hpf, for a total of 40 biological samples. Embryos were homogenized in TRIzol (Ambien). Supernatants were purified on Direct-zol (Zymo) columns with in-column DNase I digestion. RNA was eluted with RNAse-free wate, quantified. Libraries were built using Illumina TruSeq stranded mRNA poly(A) kits and sequenced on Illumina HiSeq 2500. Fastq files were aligned to genome build GRCz10 using STAR version 2.4.2a RNA-seq alignment software^15^. A gene-by-sample count table was created by counting reads that overlapped exons annotated in Ensembl GRCz10 release 91, using featureCounts function in Rsubread R package^16^. Differential expression analysis was performed using negative binomial likelihood ratio test from DESeq2 R package^17^.

Whole-mount *in situ* hybridization (WISH): Published probes for *tal1^18^, gata1a^19^, gfi1b^20^, sdc2^21^* and *cmyb* ^22^were used and WISH was performed as previously described. ^23^

Transgenic zebrafish lines: Gata1:dsRed^24^, mpx:GFP^25^ and mpeg:GFP (ZIRC) lines were used to visualize erythroid, neutrophil and macrophage populations respectively. TCF:nls-mCherry^26^ and fli1:GFP^27^ lines were used to visualize stromal cells of the CHT and the vasculature respectively.

FACS: WT and mutant transgenic embryos were grown to desired stages. Non-transgenic embryos were used as a control to set fluorescence gates. Cells were dissociated in trypsin at 37°C for 30 minutes (27 hpf), 40 minutes (48 hpf) or 1 hour (72 hpf) embryos. Reactions were stopped by adding 10% FBS and CaCl_2_ to final concentration of 1 mM. Cells were filtered through 40um nylon mesh filter (Fisher 22363547) spun at 2500 RPM for 10 minutes. Cells were re-suspended in 500 μL of 0.9X PBS/5% FBS and sorted in BD FACSAria. Sorted cells were collected in 0.9XPBS/5%FBS for May Grunwald Giemsa staining^28^. Whole Kidney Marrow FACS in adults was performed as previously described^29^.

Infection assay: 1.5 and 3 OD of lab strain *E. coli* DH5 alpha-RFP were prepared as previously described ^30^ and injected in the yolk of 2 dpf zebrafish. Each larva was kept separately in a 96 well plate and survival assessed through 7 dpf. For tracking neutrophil behavior, *E. coli* DH5 alpha-RFP was injected to 3 dpf mpx:GFP embryos grown in 150 mg/l MS-222 in embryo water. Embryos were mounted in 1% low melt agarose and time-lapse video-recorded for 10 hours to track neutrophil dynamics.

Immunostaining: Immunostaining for fibronectin (1:200) (Sigma Catalog # F3648) was performed as previously utilized^31^. Immunostaining for caspase 3 (1:200) (BD Pharmingen 559565) and MF-20(2.5 μg/ul) (DSHB) were performed as previously described^32^.

Microscopy: DIC imaging of somite borders at 22 hpf −25 hpf was performed with Nikon Widefield 90i microscope with 60X/1.00NA water-immersion objective after mounting embryos in 1% low melt agarose in presence of 150 mg/l MS-222 in embryo water at 28 °C. Infection experiment videos were captured at 28 hpf with 10X objective in NikonWidefield. For immunostained samples, images were acquired with 10X (caspase 3, MF-20) and 20X (fibronectin, DAPI) objectives with Zeiss LSM 880. May Grunwald stained cells were imaged under 20X objective with Leica DM6000 B. Image files with Z stack were processed with Fiji (Image J) to build maximum intensity projections.

## Results

### Hematopoietic pathways are altered in *MZsdc2* mutants

*Sdc2* mutant alleles were generated using TALEN technology to alter sequences in exon 2, 35 codons downstream of the start codon (Supplementary Figure 1A). HRMA^33^ was used to screen for mutations (Supplemental Figure 1C). Three independent germline lines consisting of 4 bp, 7 bp, and 10 bp deletions were identified and propagated. All deletions created shifts in reading frame leading to missense amino acid sequence and early stop codon (Supplementary Figure 1B). We screened for phenotypes in mutants obtained from heterozygous crosses (zygotic mutants) and from homozygous mutant incrosses (maternal zygotic) and found that neither the zygotic or the maternal zygotic (MZ) mutants displayed any obvious phenotypes.

Both zygotic and MZ mutants grow up to be viable and fertile with no apparent phenotypic differences compared to wildtype (WT). We performed paired-end RNA sequencing on *MZsdc2* mutants and WT embryos at the following stages of development: 1 somite, 16 somite, 30 hpf, 48 hpf and 72 hpf to screen for any transcriptomic differences. We chose these time points to cover some of the major developmental events previously implicated in morpholino screens and distinct stages of hematopoiesis^34–37^. RNA-seq analysis revealed *sdc2* as the most differentially expressed transcript (Supplementary Figure 1D) and *in situ* hybridization against *sdc2* in the mutants showed no staining, indicating nonsense-mediated decay (Figure 1B-B’). Transcript levels of *sdc3* and *sdc4* were unchanged, suggesting that the other syndecans are not modulated in a compensatory response at the transcriptome level (Supplementary Figure 1E-F).

**Figure 1.**
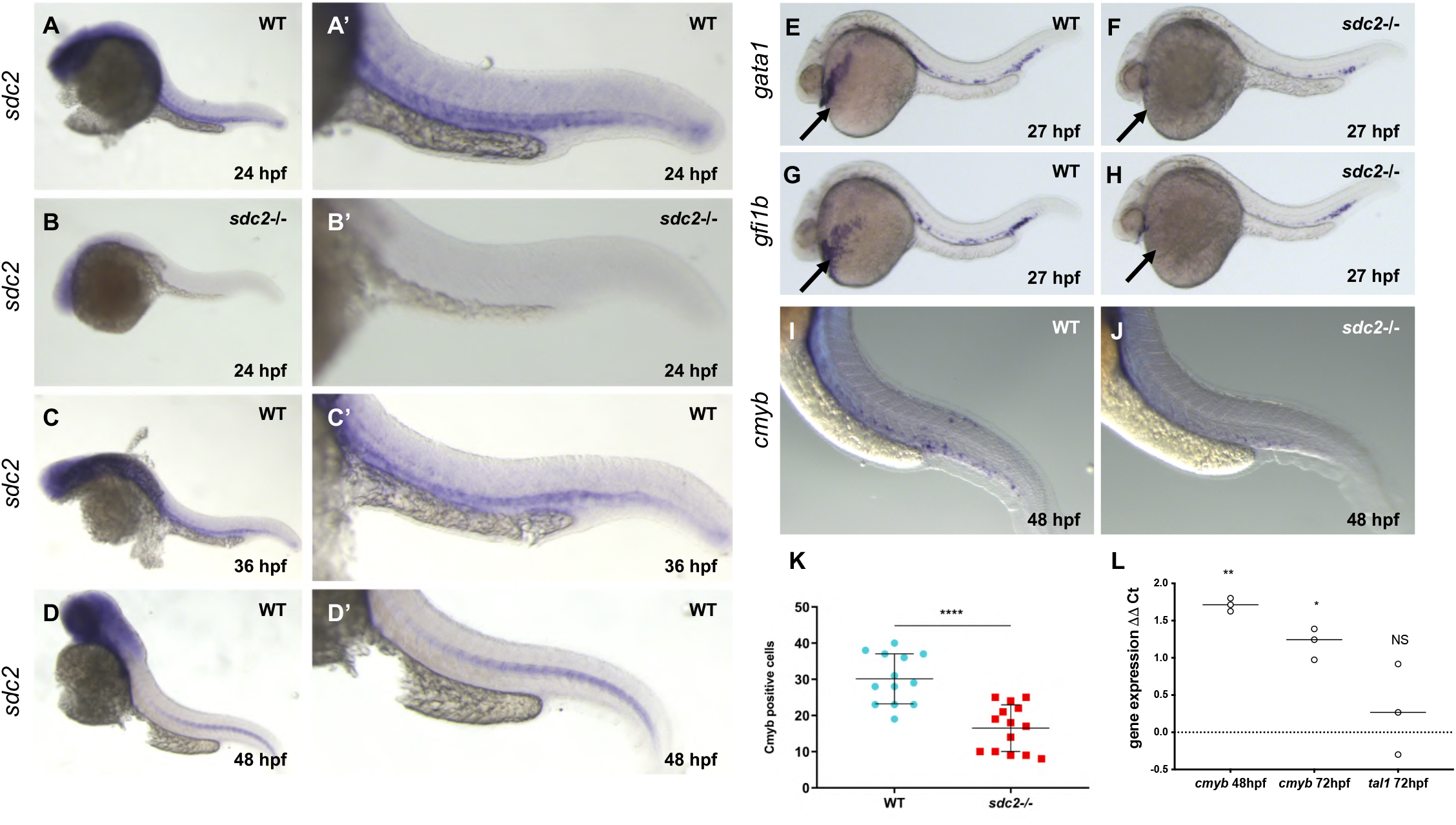
Gene expression analysis shows differential expression of hematopoietic genes in *MZsdc2* mutants. (A-A’) *In situ* for *sdc2* in 24 hpf WT embryo. (B-B’) *In situ* for *sdc2* in 24 hpf in mutant embryo showing loss of *in situ* staining. *In situ* for *sdc2* showing expression in hypochord region of WT embryos at 36 hpf (C-C’) and 48 hpf (D-D’), N=12 per genotype per time point. In *situ* for transcription factors *gata1* (E-F) and *gfi1b* (G-H) at 27 hpf. Arrow indicates loss of expression of the genes in yolk in the mutant embryos, N=20 per genotype. (I-J) *In situ* for *c-myb* at 48 hpf. (K) Quantification of *cmyb* positive cells in the WT vs mutants in the *c-myb in situ* embryos (N=13 WT, N=14 *sdc2*^-/-^). (L) Relative gene expressions of *cmyb* in mutant whole embryos at 48 hpf and 72 hpf and *scl* at 72 hpf compared to WT control. Points are individual experiments, delta Ct values used to compute standard Student’s t-test significance. NS= not significant. *In situ* images acquired with Leica DFC310 FX at 4X and 12X magnifications. Dots represent individual data points, bars represent SD, horizontal lines represent mean. Statistical significance determined using the Student’s t-test. *P<0.05; **P<0.01, ****P<0.0001.

Through differential gene expression analysis, we discovered that genes important in zebrafish hematopoiesis were differentially expressed in *MZsdc2* mutant embryos (Supplementary Figure 1G). *In situ* hybridization of *sdc2* revealed that *sdc2* is highly expressed in the hypochord region from 24-36 hpf and in the axial vasculature at 48 hpf, indicating that *sdc2* is expressed in regions of early definitive hematopoietic events (Figure 1A-D). We next analyzed expression of key hematopoietic genes, *tal1, gata1*, and *gfi1b* during primitive hematopoiesis (18 hpf) and after the onset of definitive hematopoiesis (27 hpf) by in *situ* hybridization. Expression of hemangioblast marker *tal1* was unaltered at 18 hpf and 27 hpf (Supplementary Figure 2A-D). Expression patterns of erythroid transcription factors *gata1* and *gfi1b* were normal at 18 hpf (Supplementary Figure 2E-H) but were altered at 27 hpf, absent from circulating blood and decreased in the CHT region (Figure 1E-H). To assess specification of the definitive HSPC populations, we performed *in situ* hybridization for *cmyb* at 48 hpf and qPCR for *cmyb* at 48 hpf and 72 hpf; hemangioblast gene *scl (tal1)* served as control for qPCR. Observation of *in situ* hybridization results (Figure 1I-J), quantification of *cmyb* positive cells in the hypochord region of 48 hpf WT and MZ*sdc2* embryos (Figure 1K), and qPCR for *cmyb* at 48 hpf and 72 hpf (Figure 1L) revealed that the gene expression level of HSPC gene *cmyb* was significantly lower in MZ*sdc2* at both timepoints. These results suggest that MZ*sdc2* mutants have normal primitive hematopoiesis and expression of hematopoietic markers are altered in the mutants after the start of definitive hematopoiesis, indicating that *sdc2* has a role in regulating definitive hematopoiesis.

### MZ*sdc2* mutants have immature/defective erythroid and myeloid populations after definitive hematopoiesis

Given differences in expression of some of the major hematopoietic genes, we tested whether differentiated and circulating hematopoietic populations in MZ*sdc2* mutants were altered. The *in situ* data indicated that the differences in expression of hematopoietic genes arise concurrently with the onset of blood circulation at 24 hpf. Thus, we investigated the status of erythroid cells in MZ*sdc2* mutants at various time points post 24 hpf. We pooled whole embryos at 27 hpf, 48 hpf, and 72 hpf and performed FACS using the gata1:dsRed transgenic line^38^. No change in erythroid cell count was observed in MZ*sdc2* mutants at 27 hpf (Supplementary Figure 3A-B). At 48 hpf, fewer gata1 positive erythroid cells were present in MZ*sdc2* mutants compared to WT embryos (Supplementary Figure 3C-D). In contrast, at 72 hpf there was a significant increase (p=0.004, Figure 3 C) in the total gata1 positive cells in MZ*sdc2* mutants (Figure 2A-B, red box). Further resolution of the gata1 positive population in terms of RFP intensity showed that MZ*sdc2* mutants have a significantly higher number of cells with more intense gata1 signal and a significantly lower proportion of gata1 low signal cells compared to WT (Figure 3D-E). Previous studies in zebrafish gata1:GFP transgenic lines and in mice showed that immature erythroid cells have the highest gata1 levels^39,40^. Together, these results suggest that MZ*sdc2* mutants have changes in the proportion of erythroid cells with indications of more immature erythroid cells in the mutants after the onset of definitive hematopoiesis.

**Figure 2.**
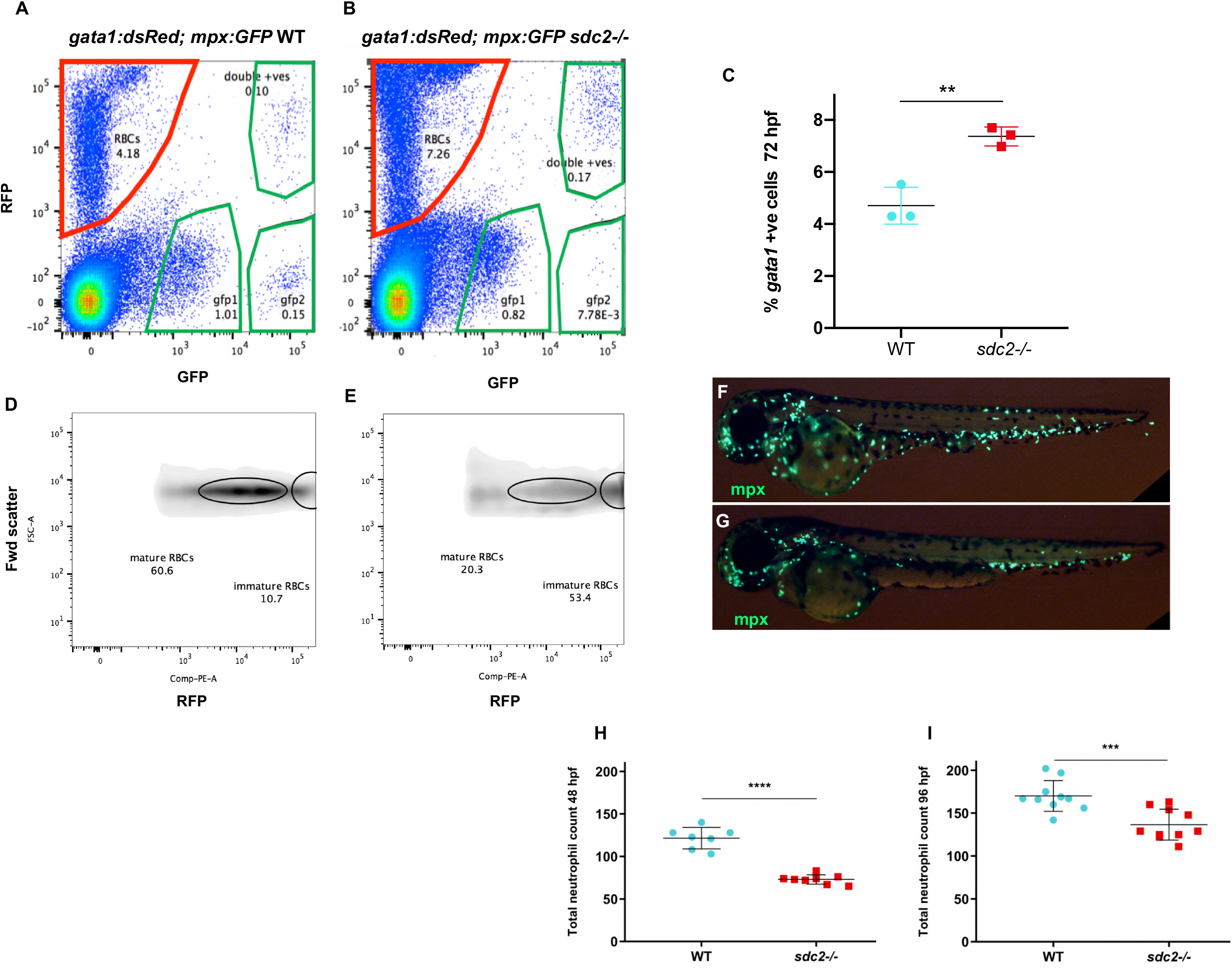
MZ*sdc2* mutants have aberrant erythroid and myeloid populations after definitive hematopoiesis. (A-B) FACS of gata1:dsRed;mpx:GFP transgenic WT and mutant embryos at 3 dpf showing the number of gata1 positive erythroid cells (y axis, red box), and mpx positive neutrophil population (green box) (C) FACS of gata1:dsRed transgenic WT and mutant embryos at 3 dpf showing percentage of gata1:dsRed positive cells in mutants compared to WT, 50 pooled embryos, per genotype, per experiment. Dots represent individual experiments. Statistical significance determined using Student’s t-test. **P<0.01. (D-E) Density plot of gata1:dsRed population at 72 hpf resolved for forward scatter in the y-axis and red fluorescence in the x-axis to better visualize the difference in RFP fluorescence intensity in WT vs mutants. Circles represent clusters of cells with high and low RFP fluorescence intensity that correspond to immature and mature red blood cells respectively. (F-G) 48 hpf mpx positive transgenic WT and mutant embryos showing the neutrophil population. (H-I) Comparison of quantification of neutrophils in the whole body of mpx:GFP positive WT and mutant embryos at 48 hpf and 96 hpf. N= 7-10 embryos per genotype, per time point. FACS performed in BD FACS Aria. Dots represent individual data points, bars represent SD, horizontal lines represent mean. Statistical significance determined using the Student’s t-test. ***P<0.001; ****P<0.0001.

**Figure 3.**
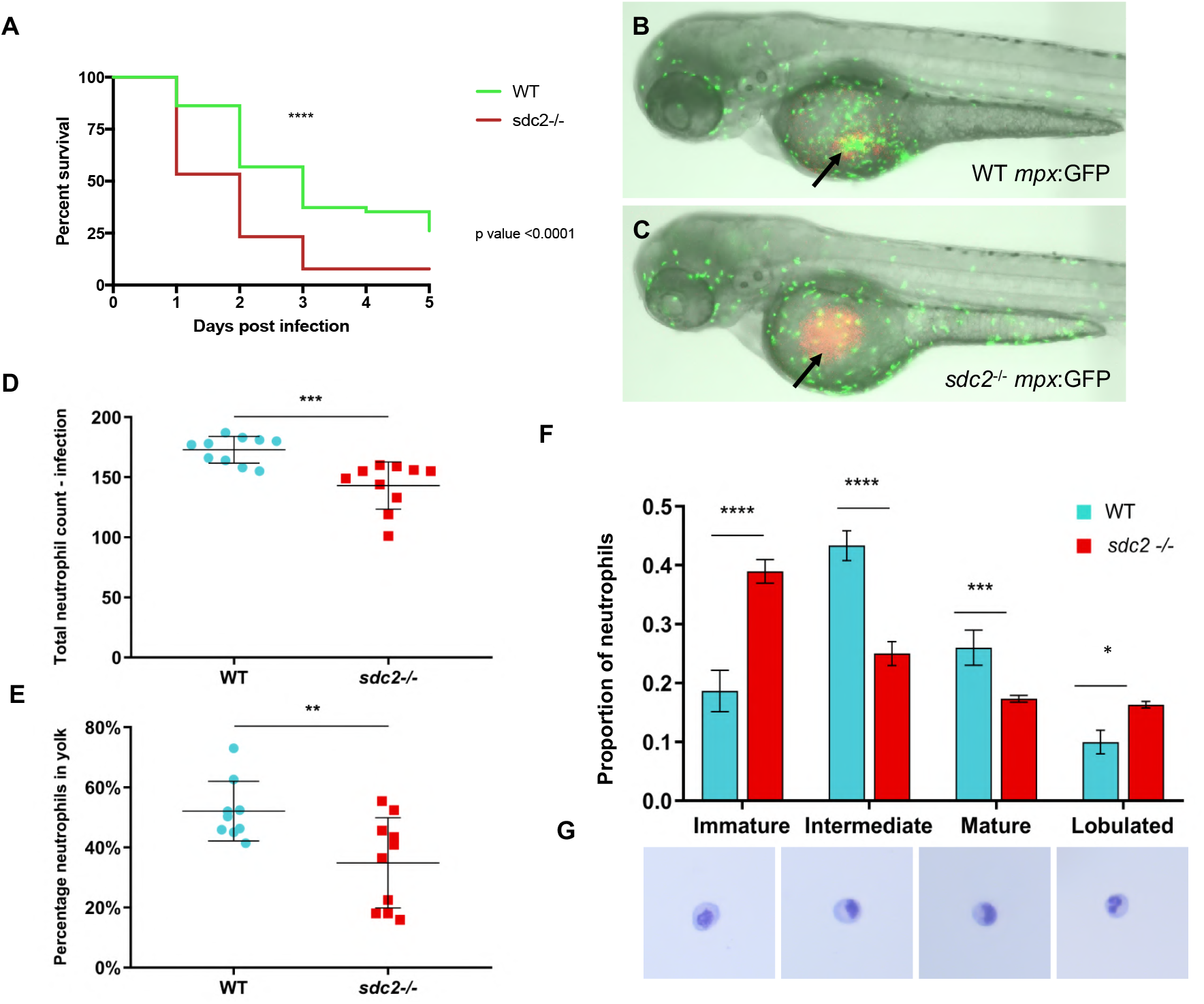
*E. coli* infection is more lethal in *MZsdc2* mutants. (A) Kaplan-Meier survival curve for WT and mutant embryos that were infected at 2 dpf and survival data was collected for 5 days post infection (dpi) (n=51 per experiment, experiment repeated 3 times). (B) 48 hpf mpx:GFP WT and mutant embryos injected with *E. coli*-RFP in the yolk. Arrows indicate site of infection in the yolk where *E. coli*-RFP was injected (N=10 per genotype). (D) Quantification of neutrophils in whole body of mutants and WT during infection. (E) The proportion of neutrophils at the yolk (number of neutrophils at the yolk divided by the total number of neutrophils in the whole body). (F) Proportion of neutrophils in different maturation categories in 3 dpf mpx:GFP positive WT and mutant pooled and FACS performed samples. (N= 50-70 embryos pooled per experiment, experiment repeated 3 times, the proportion of neutrophil at each maturation stage per experiment used to analyze t-test significance. (G) Representative images of neutrophils at different maturation stages as seen through May Grunwald Giemsa stain in WT and mutant neutrophils. Time-lapse imaging performed in Nikon Widefield at 10X objective at 28°C. May Grunwald Giemsa staining images acquired in Leica fluorescent stereoscope M205 FA with the colored camera with 20X objective. Dots represent individual data points, bars represent SD, horizontal lines represent mean. Statistical significance determined using the Student’s t-test. **P<0.01; ***P<0.001.

We next assessed the status of other two hematopoieitic populations present during early development, macrophages and neutrophils. To assess neutrophil and macrophage populations, mpx:GFP transgenes^41^ and mpeg:GFP transgenes^42^, respectively, were crossed into MZ*sdc2* lines. We followed the progression of neutrophil formation from 28 hpf-48 hpf through time-lapse of mpx:GFP embryos and found that the total number of neutrophils in the whole body was significantly lower in the MZ*sdc2* mutants at all time points.(Supplementary Video1, quantification in Supplementary Figure6). Whole-body neutrophil counts at 48 hpf (Fig 2F-H) and 96 hpf (Figure 2I) also showed that MZ*sdc2* mutants have significantly fewer neutrophils compared to WT embryos. FACS analysis of gata1:dsRed;mpx:GFP transgenic embryos at 3 dpf showed a shift in neutrophil population (green box,Figure 2A-B), with more dsRed and GFP double positive neutrophils in MZ*sdc2* compared to WT and an overall reduction in the neutrophil population.

We next assessed the macrophage population in MZ*sdc2* mutants at 3 dpf and found that MZ*sdc2* mutants had no difference in the number of macrophages compared to WT. (Supplementary Figure 4). Thus, MZ*sdc2* mutants have decreased numbers of mature erythroid cells and decreased neutrophil populations but unaltered macrophage numbers. This is consistent with previous observations that macrophages that are born in the yolk sac are present up to 96 hpf and colonize the CHT before definitive hematopoiesis commences^14^.

We next asked whether the hematopoietic populations in adults were affected in *sdc2* mutants. In zebrafish, the kidney marrow (KM) is the site of adult hematopoiesis - analogous to mammalian bone marrow^43^. We extracted KM of 6-month old WT and MZ*sdc2* mutants to analyze the hematopoietic populations present in adults through FACS. Consistent with embryonic phenotypes, alterations in hematopoietic populations were present in mutant adults compared to WT (Supplementary Figure 5A-B). Analysis of peripheral blood in mutants revealed the presence of more blast-like cells in the blood compared to the WT (Supplementary Figure 5C). Thus, MZ*sdc2* mutants have altered hematopoietic populations in the adult marrow and changes in circulating blood populations.

### *MZsdc2* mutants have altered response to infection

Given that MZ*sdc2* mutants have significantly fewer neutrophils (i.e. neutropenia^44^) and since neutropenia in clinical settings increases patients’ risk of bacterial infections, we tested whether MZ*sdc2* mutants were more susceptible to bacterial infection. We injected the yolk of 2 dpf mutant and WT embryos with two different doses (3 OD and 1.5 OD) of the DH5 alpha strain of *E. coli* that expressed RFP and recorded survival data for 5 days post-infection. *MZsdc2* mutants had significantly higher mortality, with more than 50% of the mutants dead by 24 hours post-injection (hpi) (Figure 3A).

We utilized live imaging in 3 dpf mpx:GFP transgenics to track the dynamics of neutrophil behavior in response to *E. coli-RFP* injection for 10 hpi (Figure 3B-C, Supplemental Video 2). Quantification of total neutrophil numbers during infection revealed that MZ*sdc2* mutants have significantly fewer neutrophils present in the whole body even after infection (Figure 3D).

Neutrophils accumulated at the injection site in WT embryos whereas in mutants, the number of neutrophils localized at the infection site was significantly reduced (p=0.0013, N=10 per genotype, data not shown). We next normalized the total number of neutrophils at the yolk to the total number of neutrophils in whole body for individual embryos (Figure 3E) and found that MZ*sdc2* mutants had a significantly lower proportion of neutrophils present at the infection site compared to the number present in the whole body. These results indicate that mutant neutrophils are less responsive to infection. We next performed FACS in 3 dpf mpx:GFP mutant and WT embryos, followed by May Grunwald Giemsa staining to assess neutrophil maturation. MZ*sdc2* mutants had a higher proportion of immature neutrophils and a significantly lower proportion of intermediate and mature neutrophils (Figure 3F-G). This concurred with shifts in neutrophil populations in FACS analysis of gata1:dsRed; mpx:GFP transgenic MZ*sdc2* mutants (Figure 2B). Thus, MZ*sdc2* mutants are neutropenic with immature neutrophils that are not effectively recruited to infection sites. Since erythroid cells are also defective in MZ*sdc2* mutants, we next focused on the niche in which hematopoietic stem cells differentiate.

### Ventral somite cells that give rise to the CHT niche undergo apoptosis in MZ*sdc2* mutants

*sdc2* is expressed in the ventral somite region that undergoes epithelial to mesenchymal transition (EMT) to give rise to the stromal progenitors of the CHT and axial vasculature^45,46^. To investigate whether EMT is compromised in MZ*sdc2* mutants, we performed DIC microscopy to image the behavior of cells in the ventral region of somites. Still images and videos taken from 22 to 25 hpf MZ*sdc2* mutants show many cells with altered morphology and blebbing (Supplemental video 3). By 25 hpf, mutant blebbing cells accumulated and retained a more rounded morphology compared to WT (Figure 4A-B). To test whether the accumulating cell blebs in MZ*sdc2* mutants were apoptotic, we performed caspase-3 staining of 22 hpf and 25 hpf embryos. MZs*dc2* mutants had a significantly higher number of caspase 3 positive cells in their ventral somite region compared to WT siblings at both time points (Figure 4 D-I).

**Figure 4.**
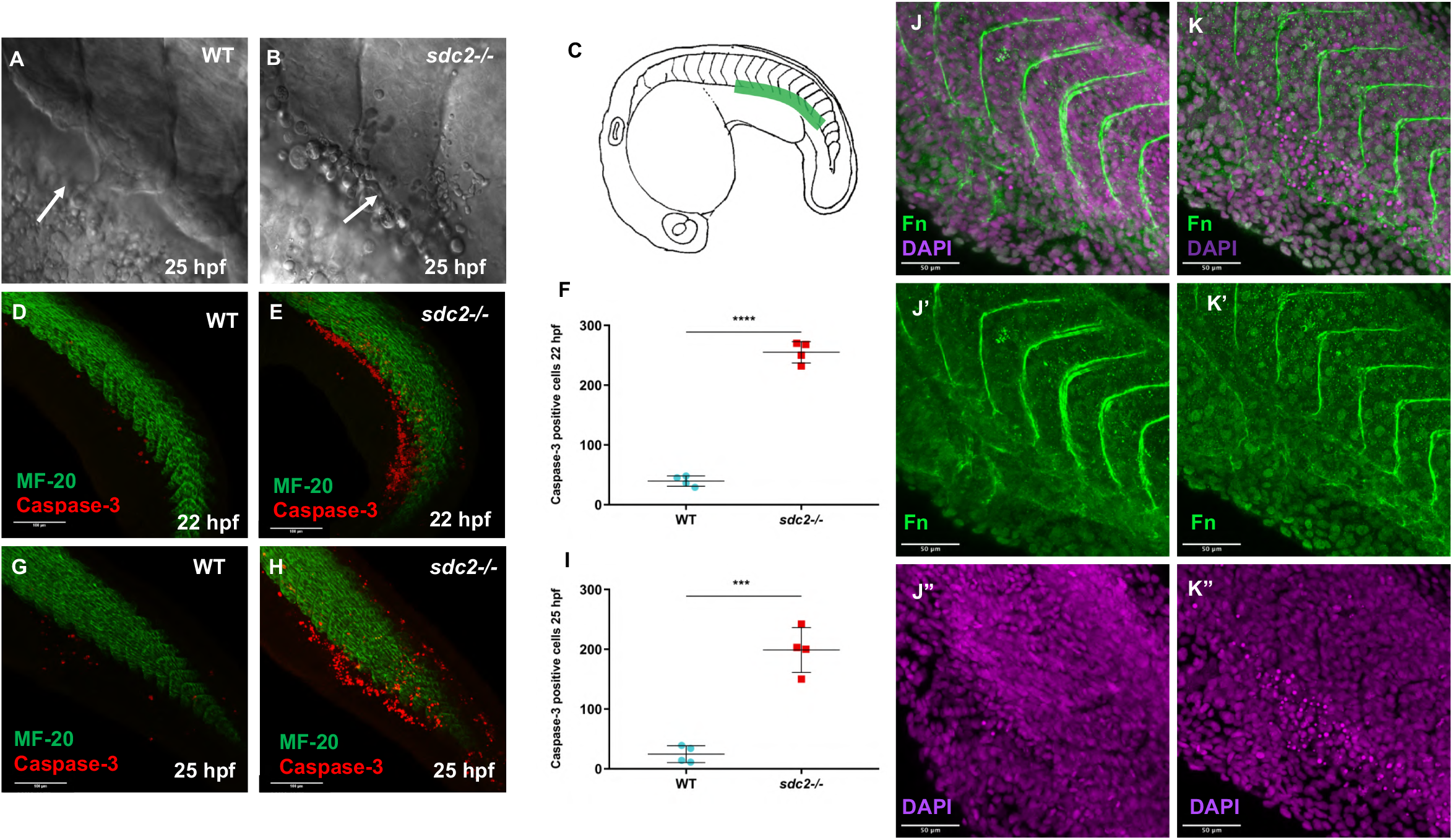
MZ*sdc2* mutants have high apoptosis in the ventral region of the somites. (A-C) DIC imaging of the ventral region of the somite at 25 hpf, area imaged indicated by green line (C). (A) Arrow indicates a cell in the process of budding off in WT and (B) accumulation of cell blebs in mutant (N=9 embryos per genotype). Caspase-3 immunostaining in WT and mutant embryos at 22 hpf (D-F) and at 25 hpf (G-I) (N=4 embryos per time point per genotype). (J-K, J’-K’) Fibronectin immunostaining in WT and mutant embryos at 22 hpf. (J”-K”) Apoptotic cells also visible as brighter cells in the DAPI channel (N=4 per genotype per time point). DIC images acquired in Nikon Widefield with 60X water lens. Immunostained samples imaged with 10X objective (caspase 3/MF-20 samples) and 20X objective (fibronectin samples) in Zeiss Airyscan. Images processed using maximum intensity projection in Fiji (Image J). Dots represent individual data points, bars represent SD, horizontal lines represent mean. Statistical significance determined using the Student’s t-test. ***P<0.001; ****P<0.0001.

The ectopic blebs were reminiscent of cells undergoing anoikis in morphants of *itga5* that have defects in fibronectin assembly^31,47^. To test whether MZ*sdc2* mutant embryos had defects in fibronectin assembly, we performed immunostaining against fibronectin in mutant and WT embryos. *MZsdc2* mutants had higher levels of unassembled fibronectin monomers demonstrated by the increased intensity of fibronectin staining all over the somite region and thinner fibronectin fibrils in somite boundaries compared to control WT embryos (Figure 4 J-K). Thus, MZ*sdc2* mutants have apoptotic cells in the hypochord region at the onset of definitive hematopoiesis, and a corresponding dysfunction in fibronectin fibrillogenesis, suggesting that loss of *sdc2* results in aberrant programmed cell death by anoikis. We predicted that the apoptotic cells in MZsdc2 mutants consist of both cell types prevalent in the hypochord region at 22 hpf and 25 hpf, the somite and vasculature cells.

### Hematopoietic niche is defective in MZ*sdc2* mutants

To test whether the apoptotic phenotype in mutants affected the CHT architecture, WT and MZ*sdc2* mutant lines were crossed into tcf:nls-mcherry;fli1:gfp transgenic backgrounds, to label CHT stromal cells and vascular endothelial cells, respectively^26,48^. Quantification of tcf:mcherry positive stromal cells in the CHT revealed that MZ*sdc2* mutants had significantly fewer mcherry positive cells in the niche at 48 hpf (Figure 5D-F). We next measured vascularization of the CHT as well as the thickness of the developing CV plexus. MZS*dc2* mutant CHT had altered vasculature with significantly fewer intervascular spaces with larger and flatter spacing than those in the WT embryos (Figure 5G-I). The thickness of the CHT however, was not significantly different (Figure 5A-C). Thus, the downstream effects of apoptotic loss of somite-derived cells and vascular endothelial cells appears to give rise to defects in the cells of the CHT potentially contributing to perturbations in HSPC maintenance, expansion and differentiation. A similar observation was made in zebrafish *olaca* mutants where apoptosis of stromal cells in the CHT leads to a defective niche that drives HSC maintenance and differentiation defects ^46^.

**Figure 5.**
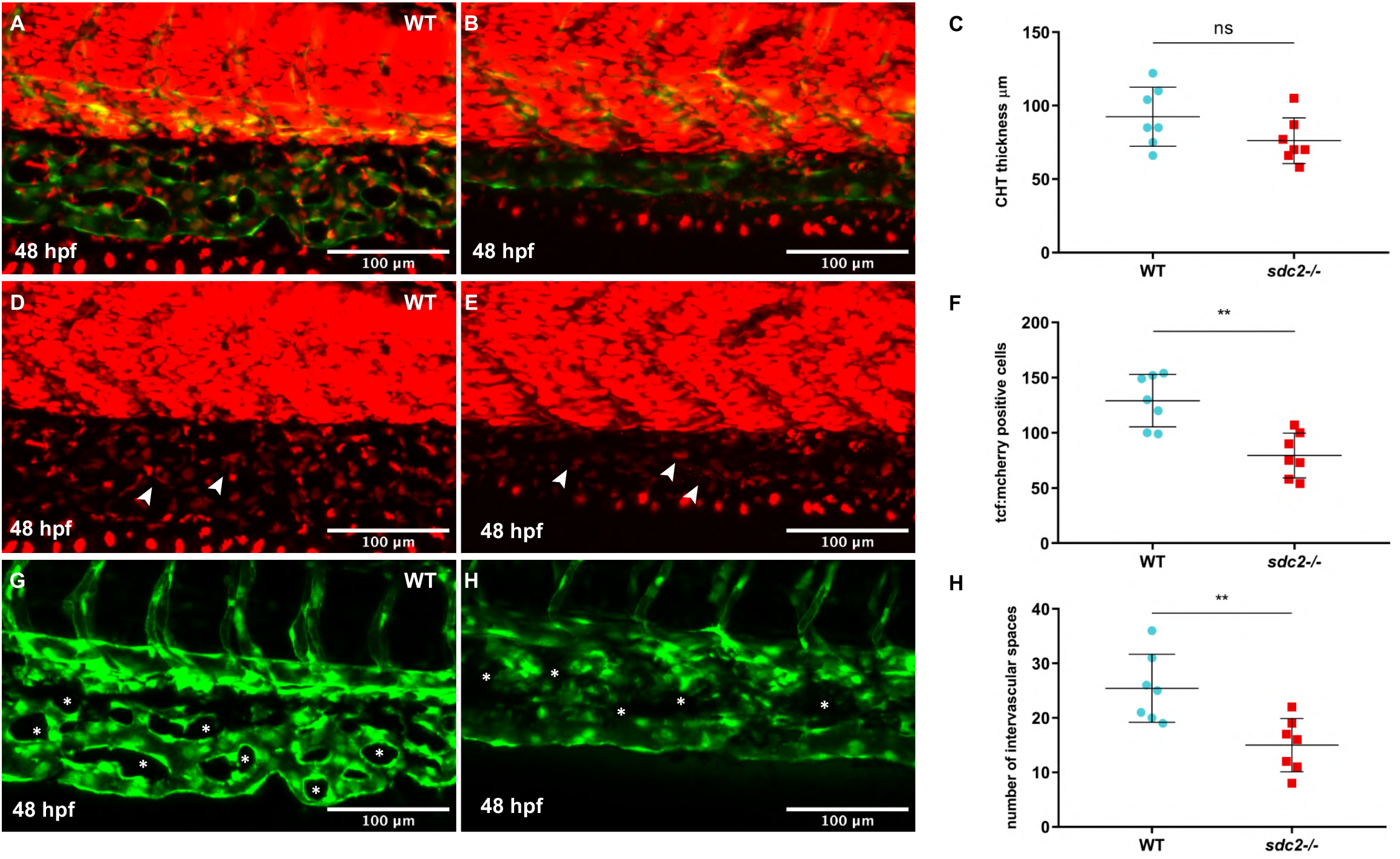
Hematopoietic niche is defective in MZ*sdc2* mutants. (A-C) Quantification of CHT thickness measurement of WT and mutant larvae at 48 hpf. (D-F) Quantification of tcf:mcherry positive cells in the CHT of 48 hpf WT and mutant larvae. Arrows represent some of the stromal cells present in the CHT. (G-I) Comparison of the number of intervascular spaces in WT and mutant CHT. Asterisks represent some of the intervascular spaces within the CV plexus. Notice that the mutant CHT has larger, flatter and less pronounced intervascular space. (N=7-8 embryos per genotype). Images acquired in Zeiss LSM 880. Images were processed using maximum intensity projection in FIJI (Image J). Dots represent individual data points, bars represent SD, horizontal lines represent mean. Statistical significance determined using the Student’s t-test. **P<0.01; NS = not significant.

## Discussion

Syndecans have been implicated in a variety of developmental and disease processes. Nevertheless, we have only scratched the surface in terms of understanding their functions in vertebrates. Our genetic analysis of zebrafish *sdc2* revealed roles in the earliest steps of hematopoiesis. We show that MZ*sdc2* mutants have alterations in the erythroid and neutrophil populations, increased mortality to bacterial infection, loss of HSPCs, and compromised CHT architecture. Our findings also indicate that the hematopoietic defects persist in adulthood.

Zebrafish is a powerful vertebrate model system to study developmental hematopoiesis as the molecular mechanisms and developmental processes of higher vertebrates are well conserved^49–51^. By virtue of studies in zebrafish^52–57^, many genes and pathways that regulate hematopoiesis have been identified. However, questions remain in understanding HSPC induction, niche environment, and factors that help differentiation and maintenance of HSPCs during early development and throughout life. Understanding these phenomena is crucial in improving HSC transplantation-based therapies.

The zebrafish CHT is equivalent to mammalian fetal liver and the first site where HSPCs home, expand and differentiate. CHT has a complex architecture consisting of stromal cells, vascular endothelial cells, and hematopoietic cells. Both the stromal cells and the vascular cells are crucial in forming a functional hematopoietic niche^14,46,58,59^. Here, we have shown that the lack of Sdc2 causes defects in stromal and vascular cells and subsequent defects in the specification of hematopoietic population. Our results suggest that the mechanism behind compromised CHT architecture in MZ*sdc2* mutants is the disruption of fibronectin assembly, leading to abnormal cell blebbing and anoikis of cells in the hypochord region. Integrin alpha-5 morphants have similar phenotypes, including disruption of fibronectin assembly and ectopic cell blebbing^31^. Besides increased anoikis due to loss of fibrillogenesis, fibronectin accumulation has been directly implicated in HSPC colonization defects in *mmp2* inhibited and morpholino studies^60^, consistent with our observations of reduced *cmyb* positive HSPCs in mutants. Previous studies in cell lines, *Xenopus* and zebrafish implicate Sdc2 in fibronectin assembly through its role as a coreceptor to integrin alpha-5^2,61^. Based on these published data and our findings presented here, we propose that Sdc2 is required to facilitate integrin-dependent fibronectin assembly for the maintenance and proper budding of the somite cells that contribute to the CHT.

In the absence of Sdc2, a failure to form a normal fibronectin matrix and a misbalance of fibronectin monomers result in anoikis of the cells contributing to the CHT.

Previous studies have also shown that Notch signaling relayed by the somites is crucial in HSPC induction^62,63^. Given that we see apoptosis and defects in the somite region in the mutants, HSPC induction may also be compromised in the mutants due to somite defects. Further exploration into HSPC numbers since their first appearance around 27 hpf should provide clues into whether HSPC induction is affected. In addition to interactions with fibronectin, Sdc2 can act as a coreceptor for Wnt, FGF or Notch signaling ligands^64–66^. Wnts, FGFs, Notch^67–69^and Cxcl12b^70^ are critical for HSPC induction. Further investigation is needed to understand whether Sdc2 also contributes to the emergence of HSPCs by regulation of any of these ligands.

We have shown that Sdc2 is necessary for normal neutrophil response to infection, but it is unknown where Sdc2 functions. Human neutrophils upregulate Sdc2 during infection.^71^ Perhaps Sdc2 is utilized by neutrophils to receive inflammatory signals or to interact with ECM during movement toward the site of infection. Alternatively, Sdc2 could function on the surfaces of other cell types to establish an infection-induced gradient of chemokines that influences the direction of neutrophil movement. For example, chemokine Cxcl8 selectively binds Sdc2 in human endothelial cells^72^ and is induced in zebrafish inflammatory response^73^.

Our results indicate that hematopoietic defects in MZ*sdc2* mutants persists into adulthood. MZ*sdc2* embryos have a proportionate decrease in mature red blood cells and increase in immature cells expressing high *gata1* RNA levels. Similarly, in adults we observe more blast like cells in peripheral blood. The Knockout Mouse Project Repository^74^ website suggests that adult *Sdc2* knockout mice have a decreased red blood cell distribution width (RDW) phenotype. RDW measures the degree of variation in red blood size^75^ and decreased RDW values often indicate anemia. Thus the role of Sdc2 in erythroid differentiation is likely conserved and warrants further exploration. In zebrafish adult mutant kidney marrow, we found shifts in proportions of hematopoietic populations. Intriguingly, in adult mouse bone marrow, *Sdc2* is enriched in long term HSCs that maintain the short term repopulating HSCs and non-renewing multipotent progenitor pool^76^. Further studies in zebrafish, mouse and humans are required to understand whether adult hematopoietic defects are a consequence of embryonic defects in HSCs and whether adult Sdc2 functions are conserved.

Our work provides evidence for a new molecule that has roles in HSPC induction, differentiation, and niche formation, and points to the need to consider HSPGs in the study of hematopoiesis. Understanding the role of Sdc2 in vertebrate hematopoiesis and establishing it as a candidate gene required for normal hematopoiesis could provide new strategies for HSC transplantation in the clinic to achieve better outcomes in treating blood-related malignancies.

## Acknowledgements

This work was supported in part by UM1 HL098160 to HJY, and sequencing was supported by a core facilities support grant to CCHCM (U01HL131003), from the National Heart, Lung, and Blood Institute. The content is solely the responsibility of the authors and does not necessarily represent the official views of the National Heart, Lung, and Blood Institute or the National Institutes of Health. We thank Dr. Scott Holley and Dr. Graham Lieschke for sharing protocols, Dr. Dave Traver, Dr. Trista North for in *situ* probes and transgenic lines, Dr. Mike Redd for providing the *E.Coli-RFP* bacteria used for infection and for his insights and feedback at the beginning of this work, and Dr. Matt Rondina for comments on the manuscript. We thank HSC cores at the University of Utah for zebrafish care, microscopy and FACS support.

## CvDC Gnomex DataHub repository RNA-seq experiments

1 somite https://b2b.hci.utah.edu/gnomex/experiments/detail/297

16 somite https://b2b.hci.utah.edu/gnomex/experiments/detail/378

30 hpf and 48 hpf https://b2b.hci.utah.edu/gnomex/experiments/detail/377

72 hpf https://b2b.hci.utah.edu/gnomex/experiments/detail/418

## Study approval

Standard protocols approved by the Insititutional Animal Care and Use Committees at the University of Utah were followed to produce, grow and maintain zebrafish embryos, larvae and adults. Zebrafish adults and embryos up until 96 hpf were used for experiments.

## Authorship and conflict-of-interest statements

Contributions: BSL, BWB, YCS, HJY designed and performed experiments; BSL, BLD analyzed results; BSL, BLD made figures; BSL, HJY wrote manuscript; BSL, BWB, BLD, HJY critically edited manuscript. The authors declare no conflicts of interest.

## Notes

### Competing Interest Statement

The authors have declared no competing interest.

