## Supplemental Figures and Methods for "Syndecan 2 regulates hematopoietic lineages and infection resolution in zebrafish"

**Supplementary Figure 1. MZ*sdc2* mutants were generated using TALEN-mediated mutagenesis.** (A) Schematic for the *sdc2* gene showing the signal peptide, extracellular, transmembrane and cytoplasmic domain. The TALEN site was 105 bp from the start codon, in exon 1. (B) Three mutant alleles, containing 4,7 and 10 bp deletions were created and all of the alleles caused missense mutation with an early stop codon. (C) High-resolution melt analysis (HRMA) was used to identify the mutants. (D) Expression of *sdc2* transcript between WT and mutants in the RNA sequencing data for 1 somite, 16 somite, 30 hpf, 48 hpf, and 72 hpf pooled embryos. (p adjusted $<10^{-40}$, log2 fold-change (mutant/WT) = 4.14, overall mean normalized counts=1154.90). (E-F) The transcript level for the other syndecans, *sdc3* (p adjusted = 0.098, log2 fold-change (mutant/WT) = 0.298, overall mean normalized counts=924.66) and *sdc4* (p adjusted=0.457, log2 fold-change (mutant/WT) = -0.086, overall mean normalized counts=5315.153) in the RNA sequencing data at 1 somite, 16 somite, 30 hpf, 48 hpf and 72 hpf. (G) Volcano plot showing significant differential expression of hematopoietic genes such as *gfi1a, mpx, hbbe1.1, itga2b and mpeg1* between the mutants and WT. Green lines indicate log 2 fold change, red line indicates p-value of 0.05.

**Supplementary Figure 2. *In situ* for *tal1* at 18 hpf and 27 hpf and *gata1* and *gfi1b* at 18 hpf show no difference in expression levels of these gene at these time points.** (A-D) *In situ* for *tal1* at 18hpf and 27 hpf shows no difference in expression between WT and mutants. *In situ* for *gata1* (E-F) and *gfi1b* (G-H) at 18 hpf shows no difference in the expression of either of the genes between WT and mutants at 18 hpf.

**Supplementary Figure 3. FACS of gata1:dsRed transgenic whole embryos at 27 hpf show no difference in proportion of *erythroid* cells, and a decrease at 48 hpf**. (A-B) FACS for gata1:dsRed embryos at 27 hpf  shows no difference in proportion of erythroid cells. X-axis refers to red fluorescence measurement and Y-axis refers to side scatter, experiment repeated 3 times. (C-D) FACS for gata1:dsRed embryos at 48 hpf shows decreased proportion of erythroid cells in mutants at that time point. 50 pooled embryos, per genotype, per experiment per time point.

**Supplementary Figure 4. FACS of mpeg:GFP transgenic whole embryos at 72 hpf in WT**

**and MZ *sdc2* mutants shows no difference in macrophage numbers in mutants**. (A-C) FACS for mpeg:GFP embryos at 72 hpf shows no difference in the proportion of erythroid cells between WT and mutants.. 50 pooled embryos, per genotype, per experiment, per timepoint. Dots represent individual experiments, bars represent SD, horizontal lines represent mean. Statistical significance determined using the Student's t-test. p= 0.960, NS= Not significant.

**Supplementary Figure 5**. **MZ *sdc2* mutants have alterations in adult hematopoietic populations.** (A-B) Whole Kidney Marrow FACS in 6 month old adults shows altered distribution of erythroid, myeloid and lymphoid populations in adult mutants compared to WT. (C) Peripheral blood smear shows presence of more blast like cells in the mutant blood compared to WT.

**Supplementary Figure 6**. **MZ *sdc2* mutants have fewer neutrophils in whole body since the appearance of neutrophils at 28 hpf.** (A) Quantification of GFP positive neutrophils at every 4 hours from 28 hpf to 48 hpf in the time lapse of mpx:GFP embryos (Supplementary Video 1). Time lapse acquired in Nikon Widefield Microscope with 10X objective at 28°C. N=11 embryos per genotype. Dots represent individual experiments, bars represent SD, horizontal lines represent mean. Statistical significance determined using the Student's t-test. *$P<0.05$,***$P<0.001$.

**Supplementary Video 1.** Time lapse of mpx:GFP positive WT and MZ *sdc2* to capture neutrophils formation from 28 hpf to 48 hpf. Time lapse acquired in Nikon Widefield Microscope with 10X objective at 28°C. N=11 embryos per genotype.

**Supplementary Video 2**. Time lapse of *E. coli* infected WT and MZ*sdc2* mutants showing differential neutrophil response. *E. coli*:RFP injected into the yolk of 3 dpf WT and mutant embryos, time lapse acquired for 10 hours post injection in Nikon Widefield Microscope with 10X objective at 28°C. N=10 embryos per genotype.

**Supplementary Video 3.** DIC time lapse of  the ventral somite region of WT and MZ*sdc2* mutant embryos capturing the abnormal blebbing and accumulation of punctate looking blebs in the mutant embryos from 22.5 hpf-25 hpf. Arrows point migratory cells post blebbing in WT animals. Images acquired every 1 minute for 90 minutes in Nikon Widefield Microscope with 60X/1.00NA water-immersion objective at 28 °C.

### Supplementary Figure 1

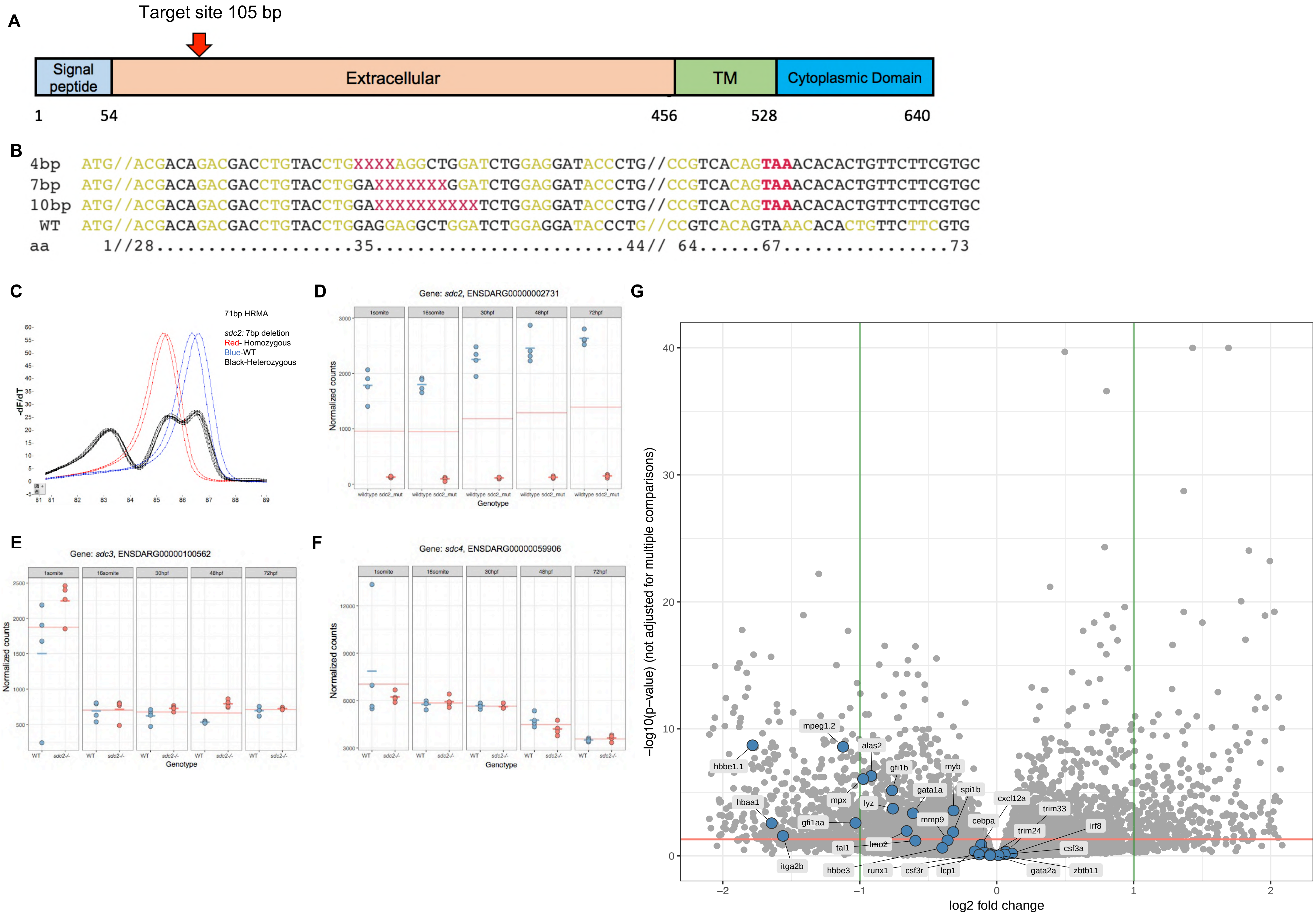

### Supplementary Figure 2

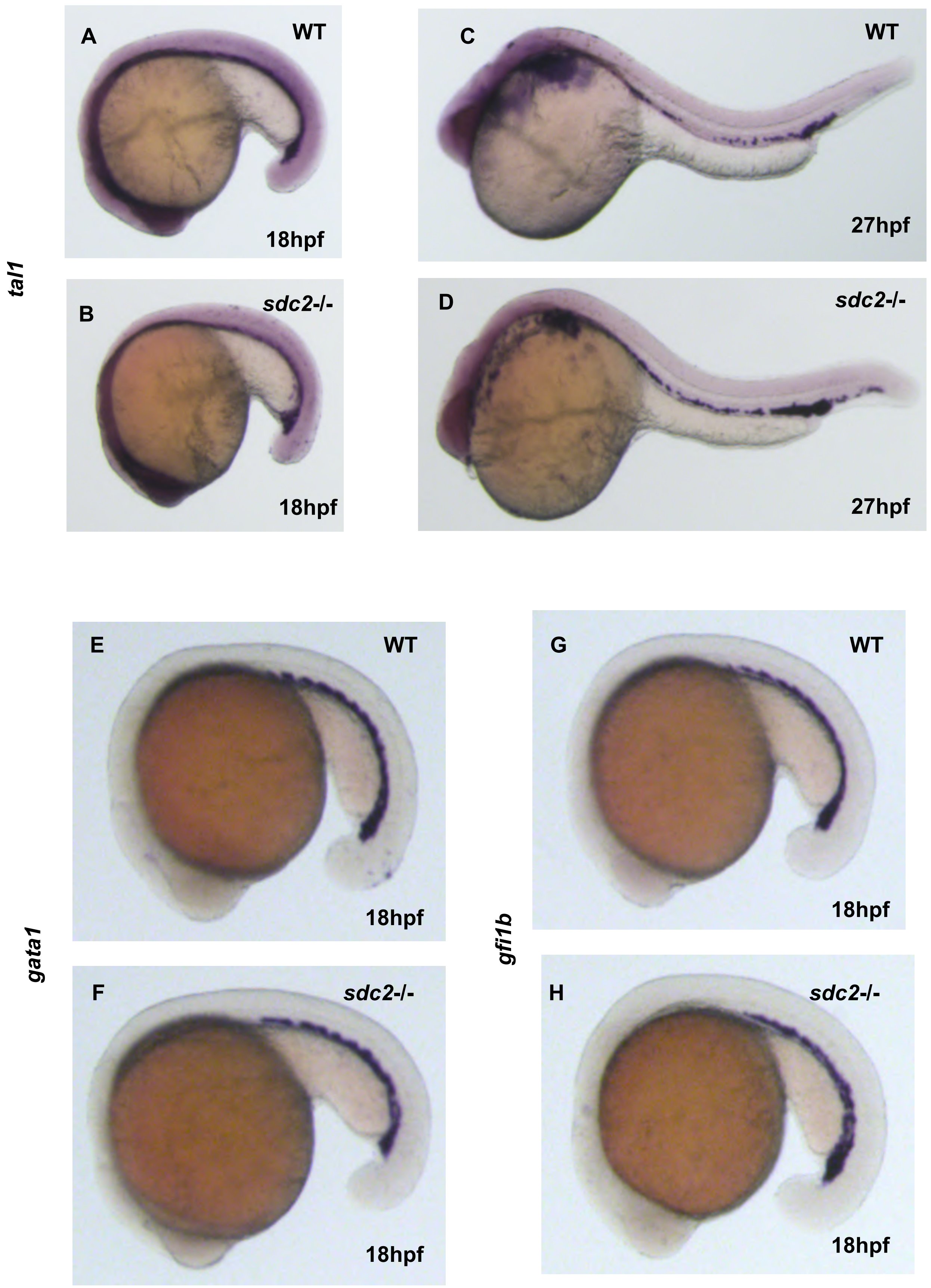

### Supplementary Figure 3

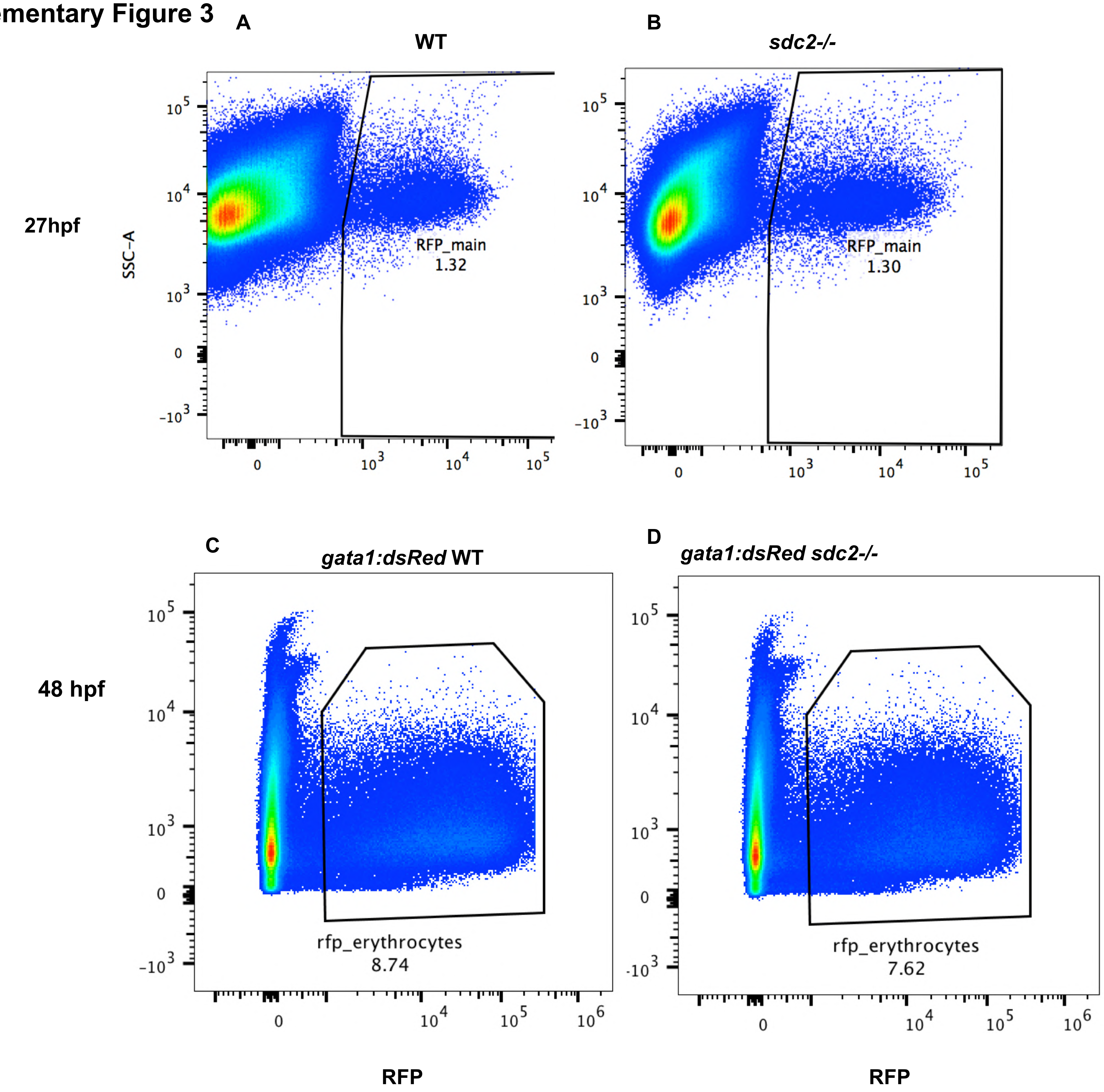

**Supplementary Figure 4**

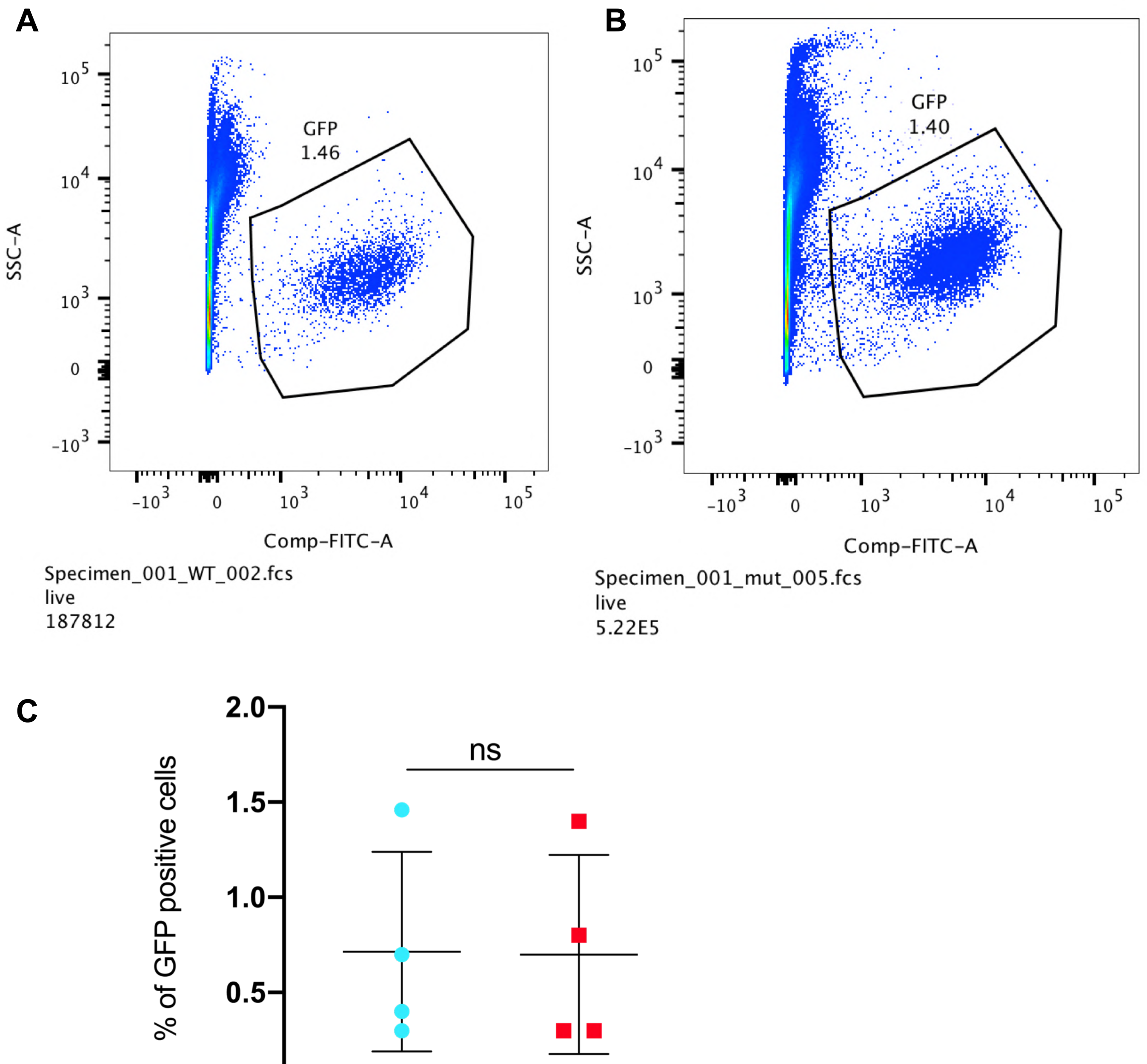

### Supplementary Figure 5

#### Adult Whole Kidney Marrow FACS

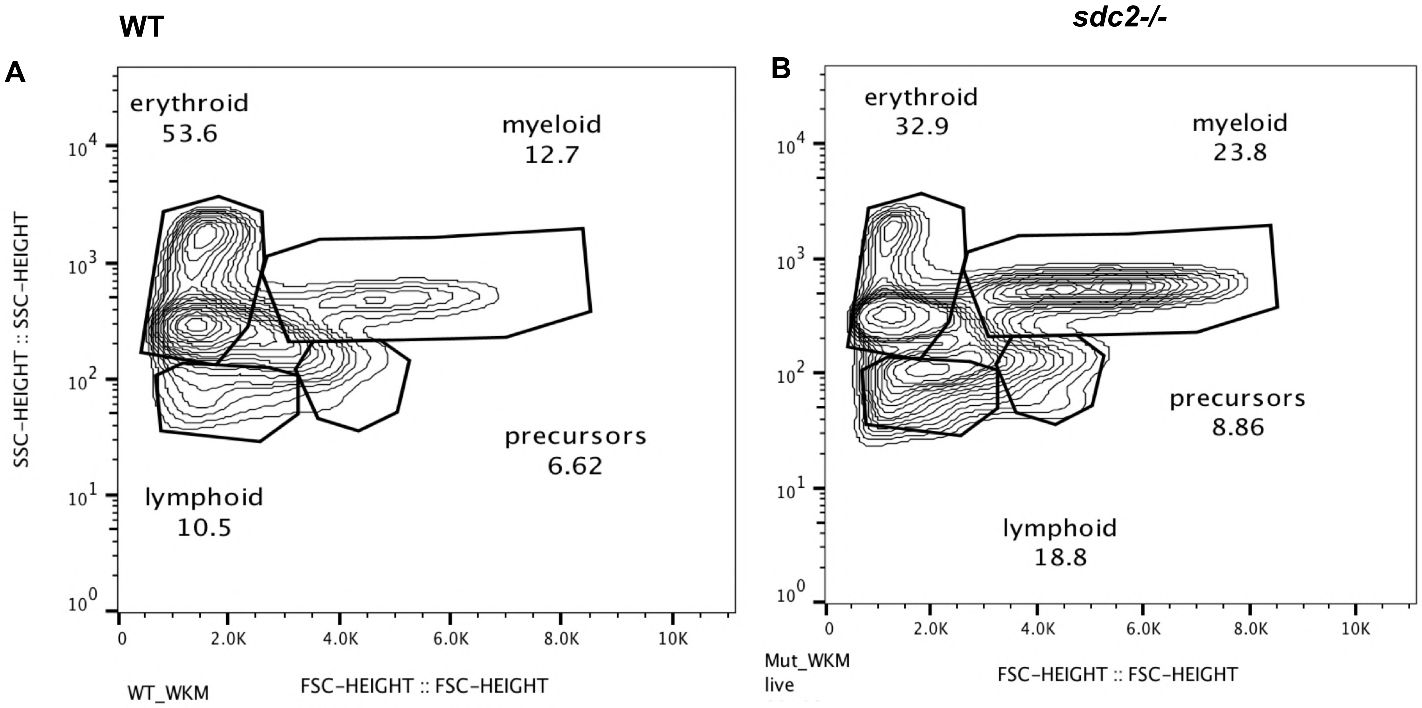


#### Adult Peripheral Blood Analysis

C

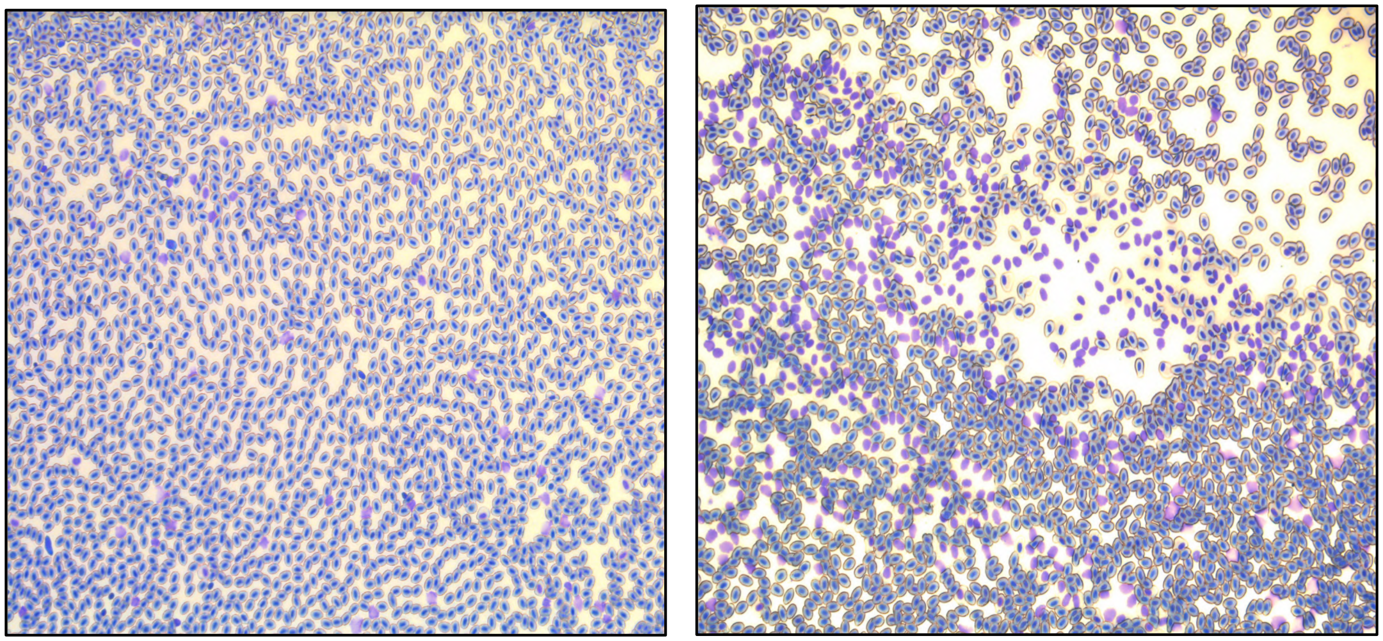

### Supplementary Figure 6

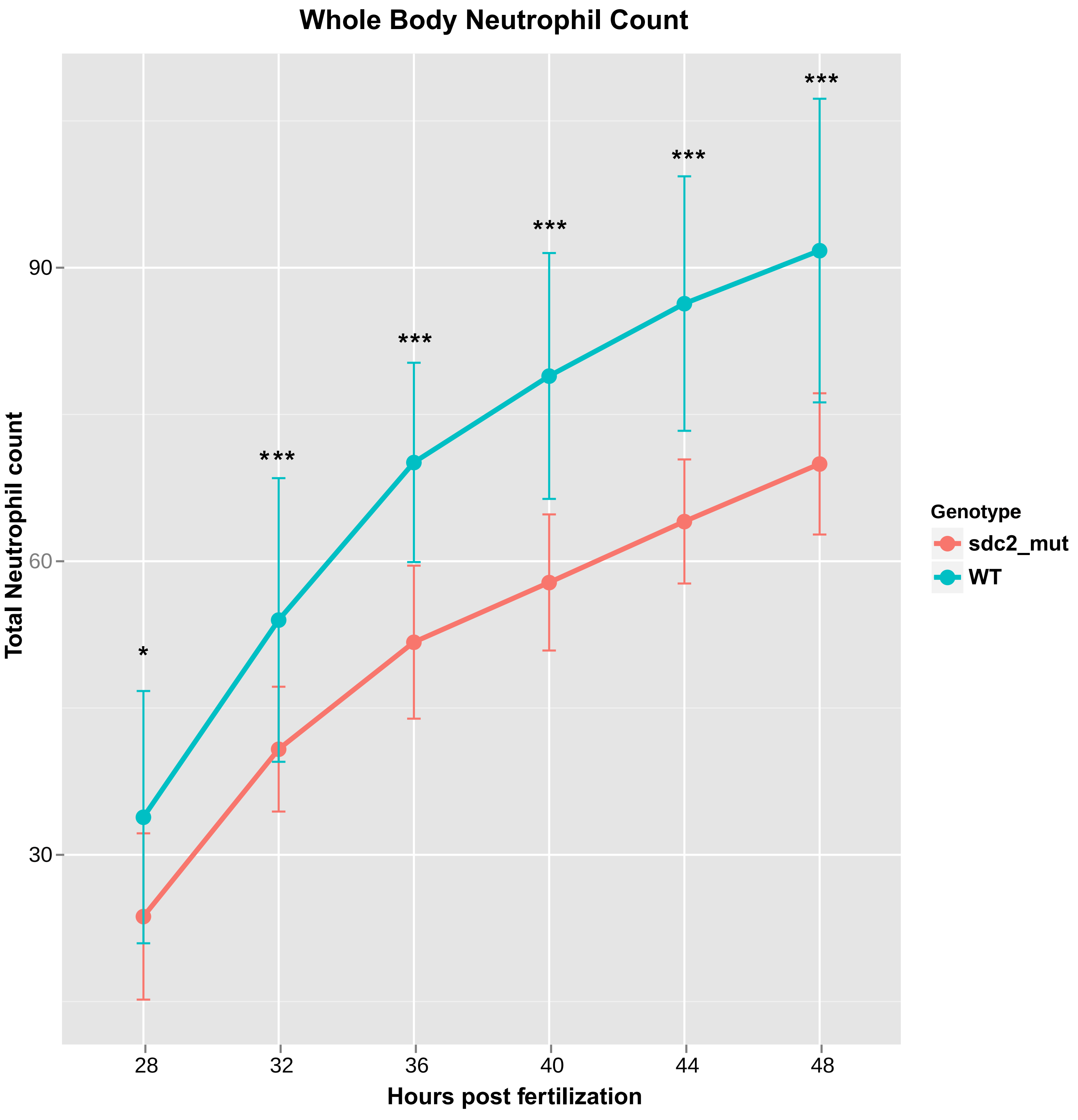

**Extended Materials and Methods**

FACS of transgenic embryos: WT and mutant embryos were grown to the desired stages. 500 μL of 30 mg/ml pronase was added to a centrifuge tube containing 50-80 embryos. The number of pooled embryos was the same between WT and MZ*sdc2* mutants for each experiment. Non-transgenic embryos were prepared for all experiments to be used as a control to set fluorescence gates. The embryos were incubated in pronase at 28.5 °C for 1-4 minutes depending on the stage then washed 3X in embryo water. 1 ml of trypsin, pre-warmed at 37°C was added to the embryos and incubated at 37°C for 30 minutes for 27 hpf embryos, 40 minutes for 48 hpf embryos and 1 hour for 72 hpf embryos. Cells were dissociated by pipetting through a 200 μL tip every 5 minutes. After complete dissociation, reactions were stopped by adding 10% FBS and $CaCl_2$ to a final concentration of 1 mM. Cells were then filtered through a 40um nylon mesh filter (Fisher Scientific 22363547) into a 50 ml falcon tube and spun at 2500 RPM for 10 min. Cells were re-suspended in 500 μL of 0.9X PBS/5% FBS (FACS buffer) and sorted in BD FACSAria. Sorted cells were collected in Trizol for RNA extraction and 0.9XPBS/5%FBS for May Grunwald Giemsa staining[43].

Cytospin and May Grunwald Giemsa staining: FACS purified mpx positive neutrophil populations and peripheral blood smear were collected in positively charged slides after cytospin at 800 r.p.m for 5 minutes. A maximum of 200 μL of cell suspension was collected per slide. Post cytospin, the slides were fixed in methanol for 1 minute. Slides were then transferred into May Grunwald solution for 5 minutes followed by 1.5 minutes in pH 7.2 PBS. The slides were then transferred to Giemsa solution (diluted at 1:10 in deionized water) for 15 minutes. Slides were then washed with deionized water and let air dry before imaging.

Whole Kidney Marrow FACS: For adult hematopoietic population, 6-month-old WT and mutant adults, (sample pooled for 3 individuals per genotype) were lethally anesthetized in 0.2% tricaine. A ventral midline incision was made to the level of the gills and the internal organs were removed with forceps. The kidney was then removed by gently teasing starting from the anterior end and working towards the posterior. The dissected kidneys were transferred to a tube containing FACS buffer. The tissue was then homogenized with a pipette and filtered over a 40 µm nylon mesh (Fisher Scientific 22363547). Cells were pelleted after adding 25 ml of FACS buffer by spinning at 280 g for 5 minutes at 4 °C. The cells were washed and centrifuged in 25 ml of FACS buffer twice and resuspended to 1ml of FACS buffer. Propidium Iodide was added at 1 µg/ml as a live/dead stain. The kidney homogenate was analyzed in Propel Labs Avalon cell sorter using Forward and Side scatter to separate hematopoietic populations in adults[44].

Immunostaining: Embryos were fixed in 4% PFA+0.2% triton overnight at 4°C. The next day, embryos were rinsed twice with PBS followed by a 3x5 minute rinse with PBS+0.2% triton. Embryos were then washed in cold acetone for 10-15 minutes (acetone step is required for caspase 3 staining only). The embryos were then washed with PBST and blocked for 1 hour in 5% goat serum in PBST (0.2% Triton). For primary antibody incubation, 2.5 µg/ul of MF-20 (DSHB) and 1:200 of caspase-3 antibody was added in 1% goat serum in PBST. The samples were incubated overnight on rocker at 4°C followed by 4 X 15 minutes in PBST on rocker at room temperature. For DAPI staining, 1:1000 DAPI was added in PBST and samples were incubated for 2-3 minutes. The embryos were then put through glycerol series with 30% glycerol in PBS for 5 minutes followed by 50% and 80% glycerol in PBS for 5 minutes before imaging.

Immunostaining for fibronectin (1:200) (Sigma Catalog # F3648) was performed as provided by Scott Holley and previously utilized[28]. 22 hpf embryos were fixed in 4% ice-cold PFA/PBS overnight at 4°C. Embryos were washed 2x5 min after PFA removal with 750 ul PBSDT( 1% DMSO, 0.1% Triton X-100 in PBS). Embryos were then digested with 10µg/ml Proteinase K in PBSDT for 4 minutes at RT followed by 2x5 minute wash with PBSDT. Embryos were fixed with 4% PFA/PBS for 30 minutes at room temperature (RT) then washed 3x5 minutes with PBSDT. Blocking was performed for 2 hours in RT in 1% blocking reagent (Roche blocking reagent 11096176001; stored as 10% stock solution at -20°C and diluted to 1% with PBSDT). Once blocking was over, fibronectin (FN) antibody (Sigma F3648) was added at 1:200 dilution in 1% block in PBSDT. The sample was incubated overnight at 4°C. Next day, samples were washed 2x15 minutes with 1% block followed by 2x15 minute wash with PBSDT. Secondary antibody (anti-rabbit, alexa 488) was added at 1:200 dilution in 1% block/PBSDT and overnight incubation was done at 4 °C. The next day embryos were washed 3x10 minutes with PBSDT followed by a 2x5 minute wash with PBST (0.1% Tween in PBS). Embryos were taken through glycerol series, 30%, 50% and 80% glycerol in PBS before imaging.
